# Automatic detection of fluorescently labeled synapses in volumetric in vivo imaging data

**DOI:** 10.1101/2025.01.22.634278

**Authors:** Zhining Chen, Gabrielle I. Coste, Evan Li, Richard L. Huganir, Austin R. Graves, Adam S. Charles

## Abstract

Synapses are submicron structures that connect and enable communication between neurons. Many forms of learning are thought to be encoded by synaptic plasticity, wherein the strength of specific synapses is regulated by modulating expression of neurotransmitter receptors. For instance, regulation of AMPA-type glutamate receptors is a central mechanism controlling the formation and recollection of long-term memories. A critical step in understanding how synaptic plasticity drives behavior is thus to directly observe, i.e., image, fluorescently labeled synapses in living tissue. However, due to their small size and incredible density — with one ∼0.5 µm diameter synapse every cubic micron — accurately detecting individual synapses and segmenting each from its closely abutting neighbors is challenging. To overcome this, we trained a convolutional neural network to simultaneously detect and separate densely labeled synapses. These tools significantly increased the accuracy and scale of synapse detection, enabling segmentation of hundreds of thousands of individual synapses imaged in living mice.

## INTRODUCTION

Understanding how learning and memory are encoded as changes to complex brain circuitry is a fundamental goal of neuroscience. One widely studied model is synaptic plasticity, in which up/down-regulation of AMPA-type glutamate receptors (AMPARs) bidirectionally tunes the strength of synaptic connection between neurons^1 2 3 4^. As AMPARs are principal mediators of excitatory neurotransmission, regulation of their relative expression gives rise to long-lasting changes in neural activity, which is ultimately thought to drive the encoding of learning and memory. Learning often involves the potentiation of excitatory synapses, while the degradation of synapses is associated with various neurological disorders, including cognitive and motor dysfunctions^5 6 7^. The ability to accurately track relative AMPAR expression over time would represent a critical leap in our ability to understand how memory is encoded in the brain. However, tracking synaptic changes over time at scale requires new computational pipelines that can automatically detect and segment individual synapses within dense imaging volumes taken from living mice (Fig. 1).

**Figure 1:**
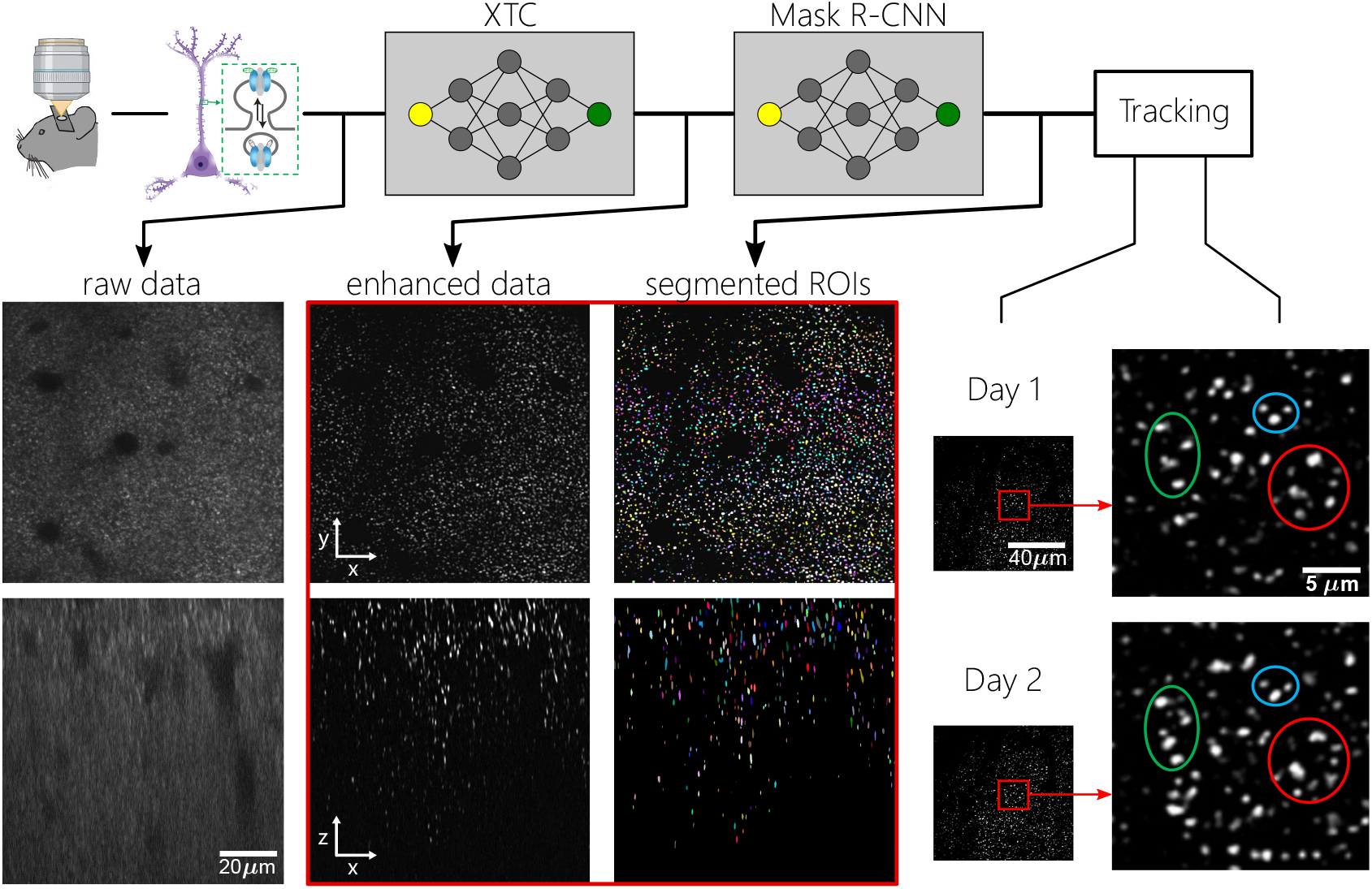
Overview of pipeline for imaging and analyzing AMPARs. Top, raw in vivo imaging data is collected, processed, and denoised using the XTC network ^9^, followed by segmentation of ROIs through MR-CNN. Segmented ROIs are then tracked across different time points to monitor synaptic changes. Images on the right demonstrate the desired goal of tracking AMPAR ROIs across multiple days, which is critically dependent on accurate segmentation.

Current cutting edge approaches to imaging synapse strength use transgenic mice to fluorescently label endogenous AMPARs, and two-photon microscopy to enable high-resolution volumetric imaging *in vivo* ^8^. Imaging these mice reveals clusters of AMPARs at excitatory synapses, where the fluorescence magnitude is proportional to the number of AMPARs^8^, a measure of synaptic strength. Two-photon microscopy is commonly used to image living tissue, as it provides superior optical penetrance and reduced diffraction, compared to confocal and other super-resolution modalities. However, current methodologies to image at micron scale in vivo are still challenged by low signal-to-noise ratios and low z-resolution due to the absence of a pinhole. To enable the annotation of individual synapses, recent work presented a computational approach to enhance image quality within such volumetric imaging datasets, called Cross-modality Trained Content-aware image restoration (XTC)^9^.

Improving the accuracy and specificity of synapse detection from large in vivo imaging datasets would substantially advance our understanding of how behavioral learning is physically manifested as precise changes in synaptic circuits. Presently, analysis of in vivo synapse imaging is conducted at the population level, describing how behavioral experience or pharmacological treatment changes the number or functional strength of all synapses in a given brain region. We seek to enhance the resolution of our understanding of synaptic encoding of behavior, focusing on tracking changes amongst millions of *individual* synapses across time.

While now possible due to image enhancement algorithms, to segment individual synapses, current approaches still require at best semi-manual annotation. The scale of in vivo synapse imaging of transgenic mice with billions of fluorescently labeled synapses makes automated detection absolutely essential, as manual or semi-manual annotation becomes time-consuming, subjective, and therefore inaccurate. However, the variability of fluorescent intensity and high density of synapses present challenges to accurately separate individual synapses from their neighbors. Synapse detection can be considered a two-step process: semantic segmentation identifies voxels that belong to any synapse; whereas instance segmentation assigns each voxel to a specific synapse, thereby disambiguating neighboring synapses. While existing algorithms such as Ilastik perform semantic segmentation of synapses with a high degree of accuracy, instance segmentation is substantially more challenging, especially in the dense synaptic networks of living brains.

Deep learning has significantly advanced the field of image segmentation, providing exceptional accuracy and efficiency in identifying and segmenting complex structures within biomedical images. Traditional image processing techniques often face difficulties with the intricate and variable nature of biological tissues, which can vary greatly in size, shape, density, and signal contrast. Deep learning models, especially convolutional neural networks (CNNs) and U-Nets, have demonstrated outstanding success in addressing these challenges because they learn hierarchical features directly from raw data. Thus, CNNs offer a promising new avenue to analyze highly complex volumetric images^10 11^. However, despite architectures such as U-Nets aving been successfully used for segmentation tasks^12,13^, such architectures were originally designed for semantic segmentation, requiring additional components for instance segmentation. Mask R-CNN is an extension of Faster R-CNN that adds a branch for predicting segmentation masks on each Region of Interest (ROI) in addition to the existing branches for classification and bounding box regression^14^. The capability of Mask R-CNN for instance segmentation is particularly suited to distinguishing individual synaptic structures from their closely abutting neighbors. Existing methods, however, are primarily tuned to the more common task in general image processing of finding small numbers of objects in complex scenes. This focus leaves a significant gap in accuracy and applicability in identifying thousands of individual objects that are densely clustered.

To achieve the goal of high fidelity automatic detection and segmentation of individual synapses, we thus adapted and trained a neural network based on the Mask R-CNN (MR-CNN) architecture. MR-CNN was trained on a new expert-annotated dataset to segment synapses in large volumetric datasets obtained from two-photon imaging in living mice. MR-CNN performs semantic and instance segmentation in parallel, predicting segmentation masks within each ROI. We demonstrate a significant increase of 20% more correctly segmented synapses—a 35.7% improvement over previous methods—in real imaging datasets, compared to separate semantic and instance segmentation using current methods. Overall, this novel network significantly enhances the accuracy and scale of synapse detection from dense fluorescent volumes imaged in living mice.

## RESULTS

We imaged fluorescently labeled synapses in living mice as previously described^9 8^. Briefly, the SEP-GluA2 transgenic mouse line expresses an AMPAR subunit fused to a pH-dependent fluorescent protein, such that functional AMPARs can be visualized when located in the neutral pH environment of the plasma membrane, but non-functional receptors located in intracellular compartments are excluded, as fluorescence is quenched at acidic pH^15 16 17 18 19^. Homozygous SEP-GluA2 mice were implanted with a cranial window and layer 1 cortex was imaged *in vivo* with a 2-photon microscope. The total volumes imaged in each stack were 98.3 µm x 98.3 µm x 60 µm, with a resolution of 0.096 µm x 0.096 µm x 1 µm. AMPAR clustering at synapses leads to punctate fluorescent signal, whose intensity is directly correlated to synapse strength^8^.

### Neural network model for combined instance and semantic segmentation of sypases

To improve the speed, accuracy, and fidelity of synapse detection, we take the approach of combining both semantic and instance segmentation into a single machine learning framework that is trained end-to-end. Specifically we adapted the Mask R-CNN (MR-CNN) architecture that has previously been designed for general image segmentation tasks^14^ to work for many similar-sized objects, instead of the original use cases of finding small numbers of the same object with highly variable size. MR-CNN is a neural network framework that can be applied to volumetric data by leveraging multi-dimensional convolutional layers, enabling us to naturally incorporate the full 3D nature of the synaptic imaging data. The core of model consists of five main sub-systems: the Backbone, Region Proposal Network, RoiAlign block, Mask head, and Classification head (Fig. 2; see Methods).

**Figure 2:**
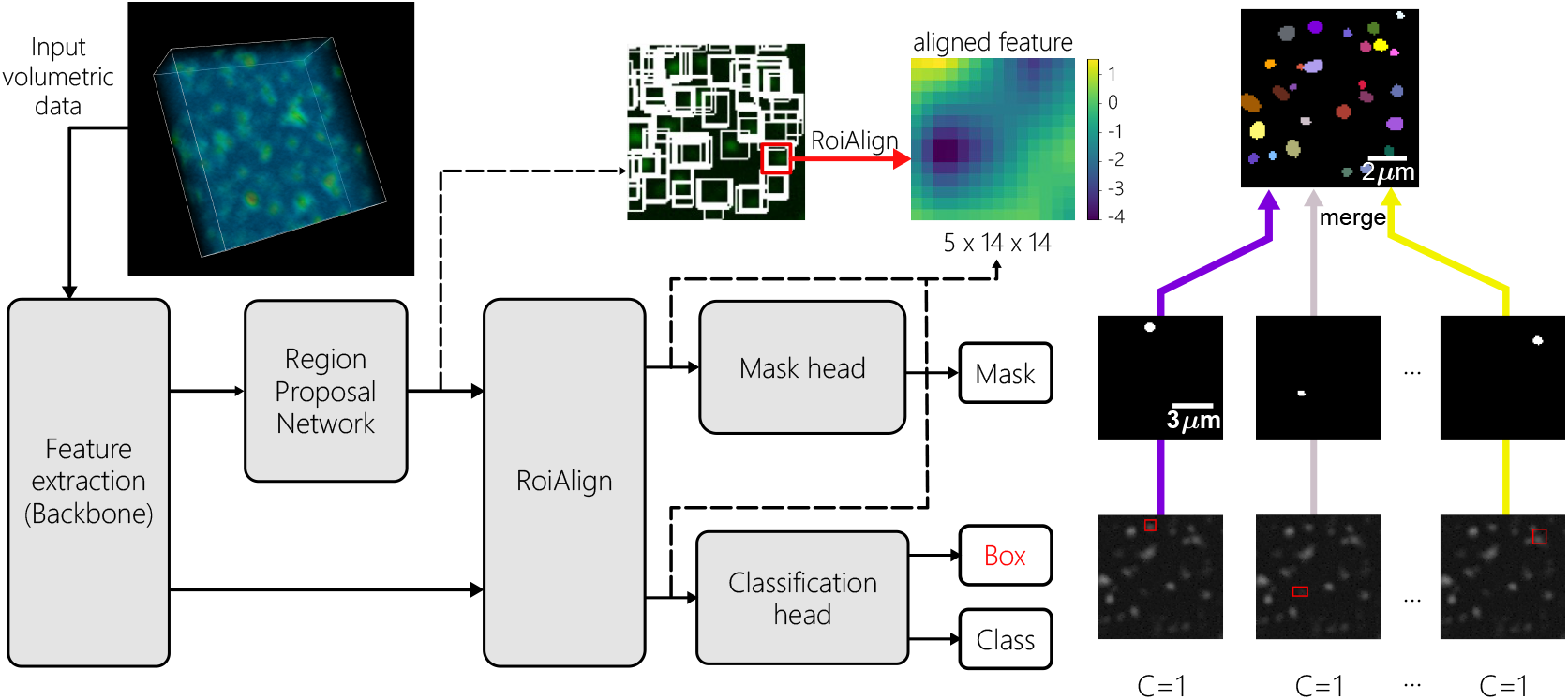
Schematic of MR-CNN architecture used for segmenting synapses in volumetric imaging data. The pipeline begins with feature extraction from the input volumetric data (top-left image) using a backbone network. Next, Region Proposal Network (RPN) generates candidate regions of interest (illustrated to the right of the input data). These regions are processed through the ROIAlign layer, which aligns the feature maps for each region (example aligned ROI shown). The aligned features are then passed to the mask head for mask prediction and the classification head for box regression and class prediction. The predicted mask, box, and class are combined to create the final segmented image of synaptic structures, as shown in the example on the far right.

The Backbone is a traditional convolutional neural network (in our case the ResNet101 architecture^20^) that provides image feature extraction. Specifically, the concatenated internal units across all layers provide a number of features at different resolutions that form the feature space in which the later networks identify the ROIs. The region proposal network then identifies, using these features, potential areas of interest as a set of bounding boxes. To facilitate the use of the backbone features within each bounding box for the final segmentation, the RoiAlign block then computes the projection of the bounding box coordinates between the feature and image space. Finally, the semantic and instance segmentations are computed on the aligned features within each bounding box by the Mask and Classification heads, respectively. The classification head computes which bounding boxes contain synapses for all potential ROIs (semantic segmentation), while the Mask head computes the specific pixels for each instance. The combination of these two produce the full segmentation.

Several key advances within this architecture were needed for efficient, accurate synaptic inference. First, many of the internal assumptions on the size and number of expected ROIs had to be modified. These changes, including changing the size ranges of bounding boxes, number of anchor points, and non-maximum suppression threshold, allowed us to incorporate the fact that synapses are numerous yet all approximately the same size. Second, the large scale of each volume prevented efficient GPU computation, requiring extensive partitioning and parallelization of the computation. While solving the memory issue, the time needed to merge the many sub-blocks was prohibitively long. We thus implemented specialized sparse-array processing that reduced the time needed for applying our model on a full volume from 12 hours to 100 minutes (a 7-times speedup). Third, to address the instability in training introduced by small batch sizes, we replaced batch normalization with group normalization (See Methods). Finally, to obtain appropriate training data, we developed a semi-manual annotation scheme (see Methods), allowing us to generate the number of samples needed to train and test the model.

### MR-CNN improves automatic synapse segmentation

In comparing MR-CNN to the previously optimized pipeline, we can analyze both the semantic segmentation, i.e., did both algorithms detect all synapse-containing voxels, as well as instance segmentation, i.e., were synapses correctly partitioned. For the former, Ilastik’s semantic segmentation captured all ROIs, meaning that semantic segmentation was not the performance bottleneck in the pipeline. Instead, the nature of Ilastik’s semantic segmentation did not ensure gaps between individual synapses, merging closely positioned synapses together. The watershed algorithm used to split these merged synapses was acceptable with significant manual checks; however, we identified two main failure modes of segmentation with Ilastik+watershed (Fig. 3). First, Ilastik combined multiple synapses and the watershed failed to detect the merge, resulting in a single instance that should have been two synapses. Second, watershed incorrectly detected a merge and split a single synapse into two, creating two instances for a single synapse. As the intensity of a synaptic punctum is directly correlated to the strength of that synapse, accurately separating individual synapses is crucial to correctly interpret the images biologically. Merging two nearby synapses into one artificially creates a false strong synapse, and the incorrect splits artificially create many weak synapses. The merging of the synapses creates giant ROIs that are not biologically realistic, and the splits likewise create small synapses that give the impression of denser synaptic connections than is correct. In addition, both merge and split errors would cause inconsistency for segmentation across days, and this inconsistency could be problematic for tracking. The two failure modes in Ilastik with watershed resulted in 45% error rate, as annotated by experts.

**Figure 3:**
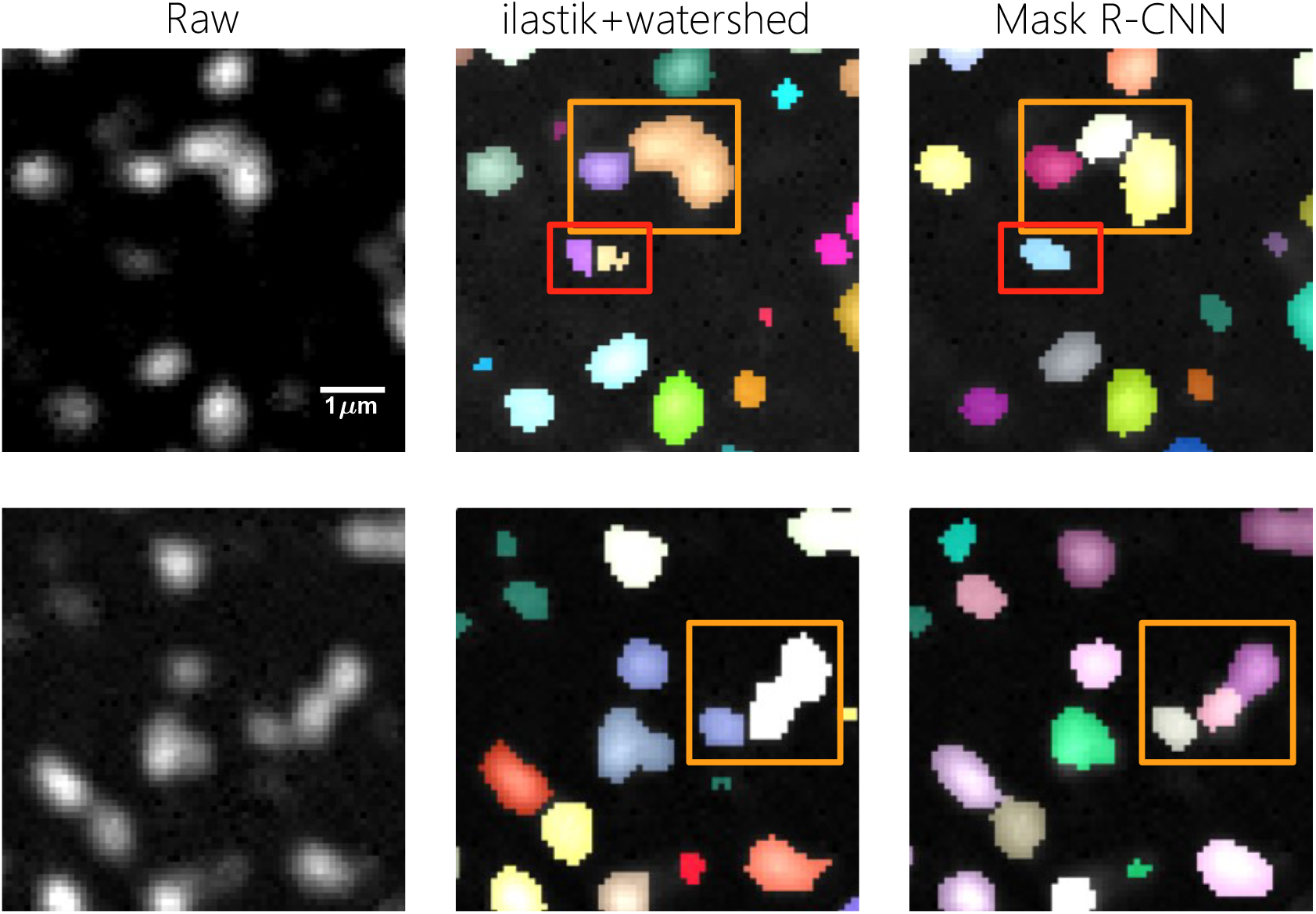
Comparison of synapse segmentation between Ilastik and MR-CNN for the same region. The leftmost column shows two examples of denoised synapse images. The middle and right columns show the same regions overlayed with segmentation results from Ilastik and MR-CNN, respectively. Individual segmentations are assigned a random color to represent the separation between each synapse. In the top row, the red box highlights an error in segmentation by Ilastik+watershed, where one synapse is incorrectly split into two. The orange boxes in both rows indicate regions where Ilastik incorrectly merged multiple synapses into one. MR-CNN provides more accurate segmentation, correctly identifying individual synapses in these regions.

**Figure 4:**
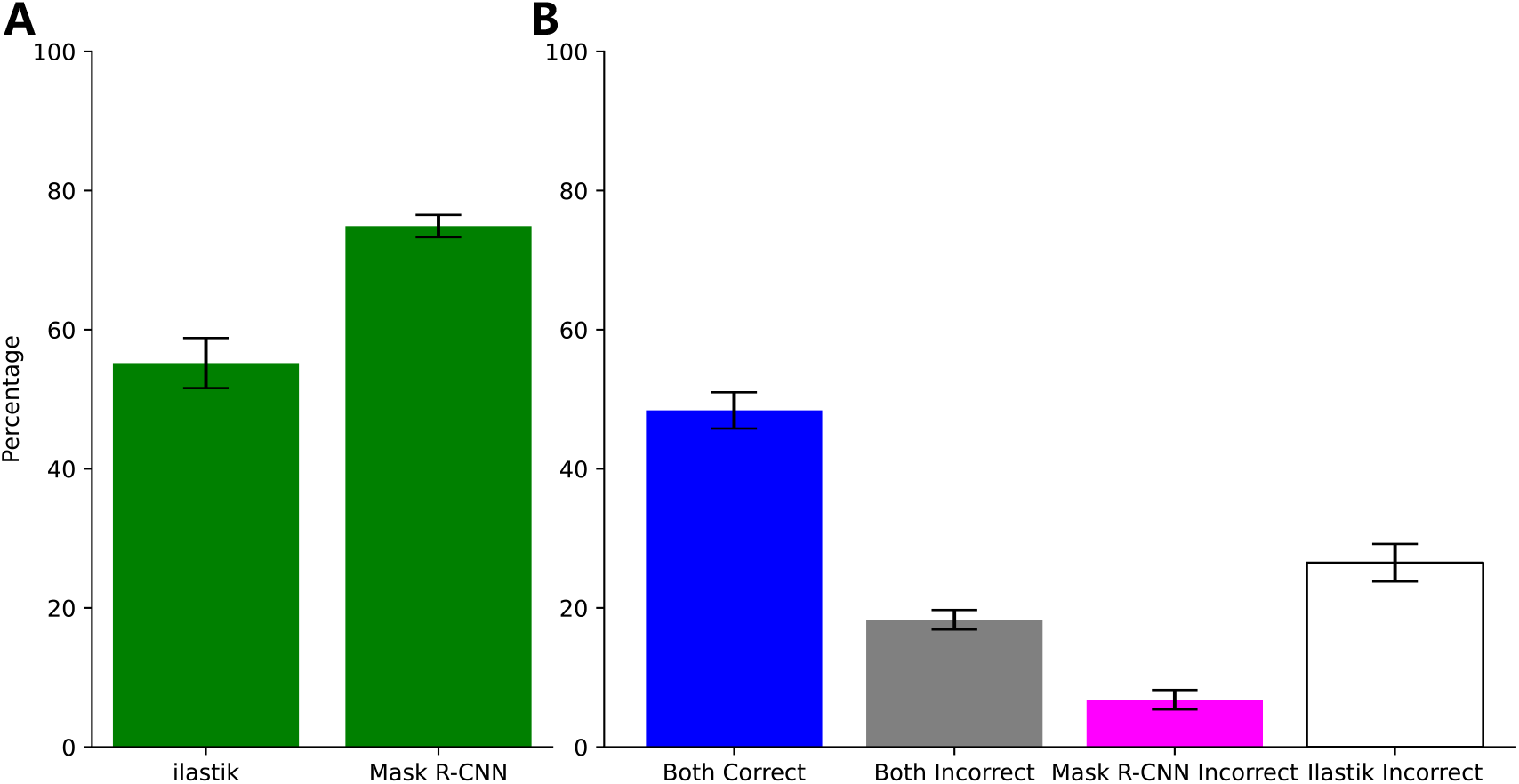
Comparison of segmentation accuracy between Ilastik and MR-CNN. A. Percentage of correct detections for both methods. MR-CNN achieved 74.9% correct detections, significantly higher than Ilastik+watershed’s 55.2% (p*<*.00001, two propotional z-test). B. Rates that MR-CNN and Ilastik are both correct (blue), both incorrect (gray), only MR-CNN was incorrect (magenta), or only Ilastik+watershed was incorrect (unfilled), with the respective percentages being 48.4%, 18.3%, 6.8%, and 26.5%. Error bars represent standard error of the mean. MR-CNN shows fewer incorrect segmentations compared to Ilastik, which has more frequent merging errors.

We trained MR-CNN on a single 1024×1024×303 volume, and we picked z-slices from 10 to 160 to avoid edge-induced artifacts. We used a 2-to-1 ratio for train versus validation (810 training and 405 validation patches). The segmentation results of MR-CNN were compared to those of both Ilastik+watershed and manual annotations. Upon manual inspection, we noticed that MR-CNN showed fewer instances of merge or split errors (Fig. 3). Specifically, we noted that in many of these examples the Ilastik+watershed pipeline was not able to correctly partition the individual synapses despite correctly identifying the same pixels at the semantic level. These examples highlighted the potential of the MR-CNN approach to more accurately segment individual synapses compared to our baseline method. Furthermore, MR-CNN performed accurate segmentation on both dense and sparse areas, avoiding false positive detections in areas where synapses are absent, such as blood vessels.

To quantify the qualitatively observed accuracy of segmentations, we required a manual assessment of both algorithms that avoided potential bias by the annotator. We thus created a graphical user interface (GUI) that allowed us to evaluate both the Ilastik+watershed and MRCNN approaches by randomly subselecting segmentations from both outputs for user assessment. The GUI first created a “review list” containing the segmentations from both methods for each ROI. The review list was shuffled, such that two segmentations on the same ROI were separated, and segmentations from both algorithms were randomly interspersed. For each segmentation, the GUI then displayed the 3D segmentation overlayed on the raw intensity image. Expert annotators then evaluated the quality of the segmentation, blinded to the algorithm that generated any particular segmentation, considering factors like segmentation accuracy, detection shape and separation from nearby synapse. Using this GUI-based annotation, two expert annotators reviewed 414 ROIs in total, from 4 different images each taken from a different mouse. On these annotations, MR-CNN demonstrated a significantly higher percentage of correct detections compared to the Ilastik+watershed approach, which had almost twice the number of merging errors (MR-CNN had only 25% error vs. Ilastik+watershed with 45% error). Overall, MR-CNN generally produced more accurate synaptic segmentation in the volumetric data. In 18.3% of the ROIs reviewed, annotators noted both algorithms created merging errors. This comparatively low level of error was expected, as previous work comparing overlap of manual detections from two expert annotators showed meaningful disagreement on the segmentation of about 15% of synapses^9^. However, while the Ilastik+watershed pipeline erroneously merged multiple synapses in an additional 26.5% of the ROIs, the trained MR-CNN was only found to make errors in 6.8% of remaining ROIs, considerably increasing its accuracy rate. The MR-CNN results are furthermore more consistent across volumes imaged in different mice, as noted by the reduced variability of more than a factor of two between images as compared to prior work (7.3% STD for Ilastik+watershed vs 3.1% STD for MR-CNN).

Finally, we complement our qualitative evaluations with additional analyses of the biological plausibility of the size of the identified synapses. This comparison considered the distribution of synapse volume across all individually segmented synapses, comparing manual annotation, Ilastik, and MR-CNN. Manual annotations were performed by annotators in a pixel-wise manner for each synapses. We compared the volume of the synapses, as these morphological characteristics reflect segmentation accuracy. Histograms of synapse volumes demonstrate that MR-CNN produced segmentations that were more closely aligned with manual annotations than Ilastik (Fig. 5). The Ilastik segmentations tended to result in larger synapse volumes (indicative of the larger number of merge errors) while MR-CNN’s results were more consistent with the manually annotated volumes. To quantify this comparison, we fit a gamma distribution—a common model for non-negative values—to the empirical distributions of the three methods. These fits reveals that the Ilastik tends to overestimate synapse volumes, producing a skewed distribution, while MR-CNN follows more closely with the manual annotations.

**Figure 5:**
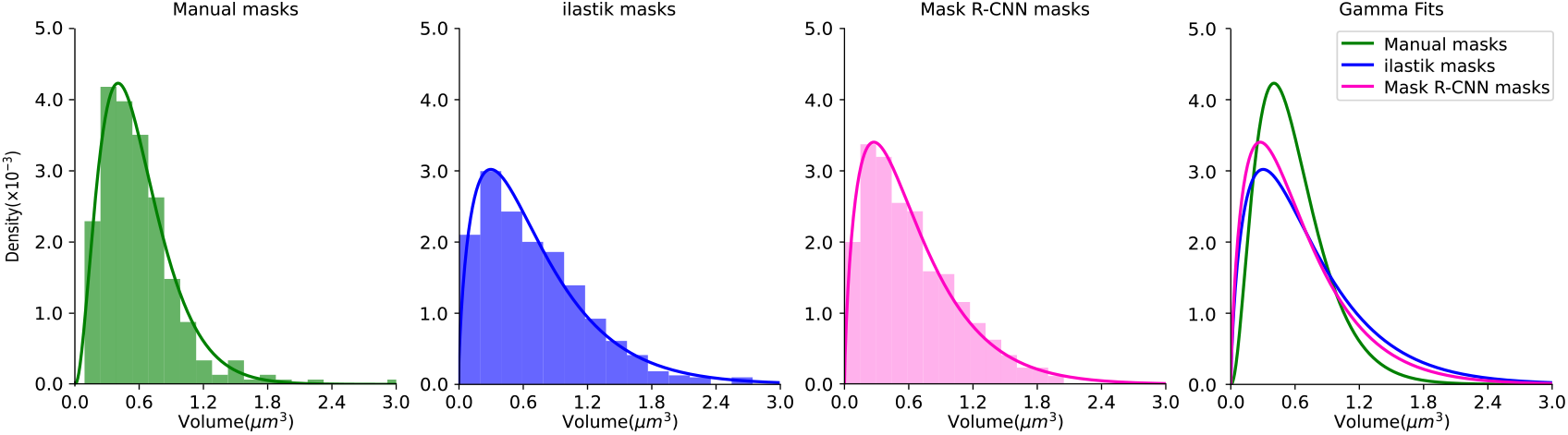
Histograms of synapse volume distributions for manual masks, Ilastik masks, and MR-CNN masks, with gamma function fits. The first three panels show the density distribution of synapse volume segmented manually, by Ilastik, and by MR-CNN. The manual annotation contains 300 synapses, while the Ilastik and MR-CNN segmentations include the same 300 synapses along with an additional 600 synapses. The final panel compares the gamma fits of all three methods, showing the differences in volume distribution across the segmentation techniques (the KL Divergence for Manual vs Ilastik, Manual vs MR-CNN, and Ilastik vs MRCNN is 0.1173, 0.0881, and 0.0117 respectively).

## DISCUSSION

This study demonstrates that MR-CNN enables accurate and robust synapse segmentation from volumetric in vivo imaging data. To optimize the model for high-density synaptic data, we modified several aspects of the network, including anchor scales and the non-maximum suppression threshold. These changes helped to accommodate the smaller size of synapses and GPU memory constraints. We used a patch-based approach, together with a merging pipeline, to resolve inconsistencies between segmented patches and minimized overlap errors. Additionally, we employed group normalization over batch normalization to address batch size limitations, improving training stability.

MR-CNN’s ability to handle instance segmentation was critical, as it accurately distinguished closely packed synapses. This is a notable improvement over Ilastik combined with the watershed algorithm, which generated significantly more incorrect merges and splits. On the other hand, MR-CNN performed instance segmentation, which was an inherent advantage for this task. Its usage of non-maximum suppression eliminated overlapping detections and separated close synapses, ultimately generating significantly more accurate synaptic detections than current methods.

In summary, the adjustments made to MR-CNN allowed it to handle synapse segmentation more effectively than Ilastik with watershed. While MR-CNN has higher computational requirements and relied on fully annotated training data, its instance segmentation capabilities make it better suited for distinguishing close synapses than semantic segmentation methods like Ilastik+watershed. Moreover, once trained, the process is fully automatic on new data, reducing the need for semi-manual annotation. The ability to label and measure the strength of every individual excitatory synapses in a volume is a significant leap in the field, refining our current population-centric view of shifts in synaptic strength. Crucially, MR-CNN improves the consistency of segmentation between mice, allowing it to be used broadly to segment similar datasets.

### Limitations of the study

There are several limitations in the MR-CNN pipeline. The model’s high computational demand restricts its scalability and can be challenging to implement without GPU resources. Additionally, MR-CNN requires fully annotated data for effective training, whereas Ilastik+watershed can function with sparse human annotations^21^. To create training data for MR-CNN, we had relied on manually curated Ilastik+watershed segmentations for ground truth. The use of Ilastik+watershed as a starting point was crucial since manually annotating synapses from scratch across large volumes is impractical. Future work could address the computational demands of the model and explore strategies for reducing the reliance on fully labeled datasets, such as weakly supervised or semi-supervised learning techniques^22^. Another limitation is the way we resolve overlapping segmentation where the patches overlap. Currently, we resolve these overlaps by selecting the largest segmentation, a choice made to optimize the efficiency of post-processing. This approach allows us to perform the selection in sparse matrix efficiently, taking approximately 100 minutes for around 330k overlapping synapses in total. However, selecting the largest segmentation is not the best strategy accounting for all overlapping instances, as it may discard smaller but valid segmentations. A more complex rule that adapts to different scenarios might be helpful in improving segmentation accuracy.

## METHODS

### Data pre-processing

Image resolution is limited by the diffraction-limited point spread function of the microscope, and imaging fluorescence from receptors expressed at endogenous levels without bleaching the signal leads to low signal-to-noise ratio (SNR). Moreover, single planes in the imaged z-stack are potentially offset due to slight movements in the the brain relative to the microscope. To improve image quality and therefore ease the identification of single synapses, we applied motion correction and image enhancement to stabilize, denoise, and sharpen the images. Specifically, we first applied pairwise affine registration on a slice-by-slice basis to align the local tissue. After registration, images were enhanced using a CNN-based denoising model built upon the Content-Aware Image Restoration (CARE) network^23^. This model, multi-modal cross-trained CARE (XTC), enhanced the SNR and resolution of the raw images to match airyscan resolution, increasing the visibility and clarity of the synapses^9^.

### Baselines and annotation

Training Mask-R-CNN requires fully annotated datasets of thousands of synapses. The size and challenging nature of AMPAR imaging prevents efficient manual annotations of the extent needed for training large-scale neural networks. Thus, we used a semi-automated approach, leveraging a previously optimized pipeline designed to analyze AMPAR imaging data^9^. The automated segment of the pipeline consisted of Ilastik for pixel-wise detection, followed by watershed algorithms to assign voxels to each individual synapses. Ilastik is a widely used interactive image segmentation toolkit^21^. Within Ilastik software, we trained a voxel-wise random forest classifier with 100 trees using the pixel classification workflow. Sparse manual annotations from expert human annotators were applied interactively to update the classifier. The voxel-wise classification performed semantic segmentation but did not assign unique labels to each ROI, leading synapses in close proximity to each other to merge into a single region. To address this, we used Matlab’s watershed function, that implements the Fernand-Meyer algorithm^24^, as a post-processing step. The watershed algorithm split “snowman-shaped” ROIs into multiple spheroid objects, and assigned unique labels to each subset of pixels. The end result of the combined Ilastik and watershed pipeline provided rudimentary instance segmentation over the entire field-of-view. These annotation were then used to train MR-CNN on the image features, and allowed us to iteratively adjust training and inference parameters to better segment the synapses.

### Network architecture

The architecture of the Mask R-CNN (MR-CNN) consists of 1) a backbone image feature extraction network, 2) a region proposal network, 3) an ROI alignment, and 4) a paired Mask and Classifiation Head (Fig. 2). The backbone of MR-CNN is a feature pyramid network (FPN), ResNet-101, that extracts feature maps from the input image. The feature maps, generated from selected layers (C2 to C5 stages) of the ResNet-101 network, are then feed into a region proposal network (RPN). The RPN produces anchor points and bounding boxes for candidate objects. The internal architecture of the RPN contains a 3×3×3 shared convolutional layer with stride of 1 to process the input feature maps. This is followed by two separate convolutional layers, each with kernel size of 1×1×1 and a stride of 1, which are used to generate anchor scores and bounding box coordinates.

ROIalign then projects the proposed bounding boxes accurately to the feature map. For each aligned bounding box, a Classifier Head predicts a class label, refines the bounding box coordinates, and a Mask Head generates a binary mask that delineates the object from the background^14^. The Classifier Head contains two convolutional layers with kernel sizes of 7×7×3 and 1×1×1, both using a stride of 1, followed by two separate fully connected layers for the class label and the refined bounding box coordinates. The Mask Head contains a series of four convolutional layers with a 3×3×3 kernel size, a stride of 1, and a padding of 1; a transposed convolutional layer with a 2×2×2 kernel size, a stride of 2, and a ReLU activation function; and final a convolutional layer with a 3×3×3 kernel size, a stride of 1, followed by a sigmoid activation function.

Due to the size of the data, and the ability of the imaging technology to further grow, memory constraints played an important part in our design. Specifically, the RPN feature maps in MR-CNN were limited to 128 to conserve GPU memory. Similarly, we used a batch size of two to stay within the constraints of the GPU memory. This small batch size introduced noise and instability when using batch normalization, so we applied group normalization to mitigate the variability in batch statistics^25^. Our switch to group normalization from batch normalization effectively compensated for the small batch size.

Traditionally, the MR-CNN architecture was used to identify small numbers of objects that could be in the foreground or background, thereby changing sizes dramatically. In our application, we did not expect synapses to be of drastically different sizes. However, we expected to have many more synapses overall per volume. Consequently, we adjusted a number of internal settings in the MR-CNN framework to appropriately address the dataset’s unique characteristics. One important alteration was to change the anchor scales — which control the potential size of objects — to [2,4,8] for each x, y and z dimensions, reflecting the relatively uniform and small size of synapses. Furthermore, the non-maximum suppression threshold for the final bounding box selection was set to 0.2 to gain sufficiently dense segmentations while attempting to minimize overlapping boxes. The latter step was critical as the sheer number of synapses in densely packed tissue could easily strain the computational resources without proper limitations.

### Training MR-CNN

The network model was trained end-to-end on the semi-automatically labeled training set. The full loss function *L* for MR-CNN is the sum of three losses:

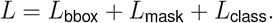

In this cost, *L*_bbox_ is the loss for the bounding box placement, *L*_mask_ is the loss for the pixel-wise mask, and *L*_class_ is the loss for the class labels per instance. The bounding box loss 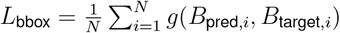 where *B*_pred,*i*_ and *B*_target,*i*_ are the *i*^*th*^ predicted and target bounding boxes, respectively, *N* is the total number of identified ROIs, and *g*(·, ·) is PyTorch’s Smooth L1 Loss, defined as:

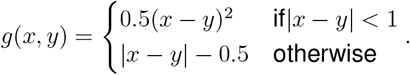

The mask loss

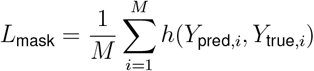

where *M* is the number of ROIs, *Y*_pred,*i*_ and *Y*_true,*i*_ are the predicted mask and the ground truth mask for ROI *i*, respectively, and *h*(*·, ·*) is PyTorch’s binary cross-entropy loss, defined as: *h*(*x, y*) = *−y* log(*x*) *−* (1 *− y*) log(1 *− x*). The classification loss

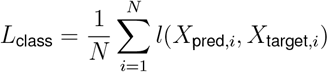

where *N* is the number of sampled ROIs, *X*_pred,*i*_ is the predicted class logits, *X*_target,*i*_ is the true class ID for ROI *i*, and *l*(*·, ·*) is Pytorch’s cross-entropy loss, defined as

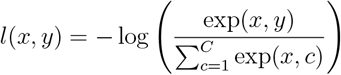

where *C* is the total number of classes.

The training images, obtained through the XTC enhancement pipeline were segmented using the semi-automated approach at the whole image scale. The segmented volume was then divided into patches for training. MR-CNN was trained using pytorch running on NVIDIA Quadro RTX 5000 GPU with 16 GB of memory. We compared training with different patch sizes, and while larger patch sizes generally performed better, GPU memory constraints limited the training patch size to 96×96×32. Patches of this size typically contained about 150 synapses with an average size of approximately 200 voxels. The number of examples per volume was especially important in providing MR-CNN both sufficient contextual information and sufficient examples per patch. Volumes that contained very few synapses were removed and the remaining were cut into patches without overlap. For every ten patches, one was turned into black background to serve as negative samples, which are examples of the background needed by the MR-CNN algorithm.

After initial rounds of training, we noticed that the trained model had bias on the direction of the image and the location of the synapses. Specifically, all pixels in the output masks were shifted by a constant number of pixels. To overcome this bias, we employed data augmentation. For each patch, we introduced four different versions with one-pixel shifts on x and y direction (i.e., x+1, x-1, y+1, y-1). Moreover, each patch has equal probability to be rotated in x, y, or z direction by either 0, 90, 180, or 270 degrees.

Once training patches were generated, every segmentation was automatically reviewed in order to remove possible false-positive detections. If a z-slice in a segmentation was either less than 3 pixels or consisted of a straight line of pixels, those pixels were removed from the segmentation. Once each slice was analyzed, all detections were relabeled in three dimensions using the python skimage.measure.label function, with a connectivity of 1, to ensure cohesive segmentations. Then, any segmentation that contained fewer than 3 z-planes were removed. These criteria improved the reliability of our training data by reducing segmentation artifacts from the Ilastik and watershed guided labeling, and removed segmentations that were almost completely cut off due to the patch creation.

### Model inference

We performed inference on images sized 1024×1024×303 not used in the training set. To efficiently manage memory, the images were split into 96×96×32 sized patches with overlaps of 34 pixels in the xy plane and 17 pixels in the z direction, ensuring all synapses remained intact within at least one patch. However, this patch-based approach led to fragmented and inconsistent detections at patch boundaries and where patches overlapped, so we applied post-processing steps to handle overlapping detections. After the network processed each patch, only bounding boxes with scores greater than 0.99 were retained. For these selected boxes, we extracted the pixel coordinates of each mask and stored them locally for parallel processing. Each mask’s coordinates were flattened into a 1D format and stored in a sparse column matrix of size 1024×1024 ×303 by 1. We concatenated all the masks into a large sparse matrix *A* of size 1024 ×1024 ×303 by the total number of masks.

To resolve the overlapping masks, we calculated the overlapping area between each mask by *A*^*T*^ *·A*. Then we used the connected components function from scipy.sparse.csgraph to identify connected components in matrix *A*. Within each connected component group, we selected the largest masks of each group, and the ones that had less than 10 pixels of overlap with the largest one were retained for the next round. After selecting and retaining recursively, the masks selected in each round were converted to a dense version and merged into one unified image. This iterative merging approach produced a single cohesive segmentation, resolving inconsistencies across overlapping patches.

## RESOURCE AVAILABILITY

### Lead contact

Requests for further information and resources should be directed to and will be fulfilled by the lead contact, Adam S. Charles.

### Materials availability

This study did not generate new materials.

### Data and code availability

- MR-CNN Code will be made available on Github github.com/zhc008/MRCNN
- XTC Code for pre-processing is available on Github github.com/yxu233/Xu_and_Graves_XTC_syn_tracking
- The Ilastik software is available at https://www.ilastik.org/

## ACKNOWLEDGMENTS

This work was funded by the National Institute of Health R01 NS134842 to R.L.H., A.R.G., and A.S.C. The authors thank all members of the Huganir, Graves, and Charles labs for their support, as well as Drs. Benjamin Pedigo and Tiger Xu for helpful discussions on efficient algorithmic implementations.

## AUTHOR CONTRIBUTIONS

Conceptualization A.S.C. and A.R.G; methodology Z.C., G.I.C., A.R.G., and A.S.C; investigation Z.C., G.I.C., E.L., A.R.G., and A.S.C.; writing original draft Z.C., G.I.C., A.R.G., and A.S.C.; editing Z.C., G.I.C., A.R.G., and A.S.C.; funding acquisition R.L.H., A.R.G., and A.S.C; resources R.L.H., A.R.G, and A.S.C; supervision A.R.G., H.I.G., and A.S.C.

